# RNA liquid biopsy via nanopore sequencing for novel biomarker discovery and cancer early detection

**DOI:** 10.1101/2025.07.02.662774

**Authors:** Vikas Peddu, Alexander Hill, Sree Lakshmi Velandi Maroli, Connor Mattingly, Joshua M.V. Gardner, Karen H. Miga, Rebecca C. Fitzgerald, Daniel H. Kim

## Abstract

Liquid biopsies detect disease noninvasively by profiling cell-free nucleic acids that are secreted into the circulation. However, existing methods exhibit low sensitivity for detecting early stages of diseases such as cancer. Here we show that long-read nanopore sequencing of full-length cell-free RNA in plasma from healthy individuals, precancerous Barrett’s esophagus patients with high-grade dysplasia, or patients with esophageal adenocarcinoma reveals a diverse cell-free RNA transcriptome that can be leveraged for detecting and treating disease. We discovered 270,679 novel, intergenic cell-free RNAs, which we used to build a custom transcriptome reference for quantification, feature selection, and machine learning to accurately classify both precancer and cancer. Moreover, we found potential therapeutic targets, including metabolic, signaling, and immune checkpoint pathways, that are highly upregulated in both precancer and cancer patients. Our findings highlight the utility of our RNA liquid biopsy platform technology for discovering and targeting early stages of disease with molecular precision.

---

Esophageal cancer is the sixth leading cause of cancer-related death around the world, with a 5-year survival rate of approximately 20%^1^. An estimated 544,100 people died from esophageal cancer in 2020, and current projections estimate that 880,000 people will die from esophageal cancer in 2040^2^, highlighting the need for better methods to detect and treat esophageal cancer at the earliest signs of disease. Gastroesophageal reflux disease can lead to precancerous Barrett’s esophagus and dysplasia, which progressively increases the risk of developing esophageal adenocarcinoma^3^. However, currently available liquid biopsy methods based on cell-free DNA sequencing detect early-stage cancers with low sensitivity and do not provide comprehensive insights into potential therapeutic targets^4^.

Liquid biopsy technologies based on cell-free RNA (cfRNA) sequencing show promise for the detection of certain cancers and precancerous conditions^5–12^. cfRNA is actively secreted by cells into the circulation and protected from degradation by being encapsulated in extracellular vesicles (EVs), which are nanometer-sized lipid bilayer particles that reflect the state of their cell of origin^13–15^. cfRNAs are comprised of various classes of short and long RNAs, including protein-coding RNAs, long noncoding RNAs (lncRNAs)^16,17^, and repeat element-derived RNAs^18–21^. Although repeat-aware RNA liquid biopsy technologies such as COMPLETE-seq^5^ enable the accurate classification of certain cancer types, including lung, liver, colorectal, and stomach cancers, other cancers such as esophageal cancer exhibit lower sensitivity, highlighting the need for novel approaches that enable cancer early detection with high sensitivity and specificity.

Current RNA liquid biopsy approaches are also constrained by the use of targeted approaches and/or short-read sequencing technologies, which are unable to sequence full-length cell-free RNAs that are longer than a few hundred nucleotides in length. Single-molecule, long-read nanopore sequencing enables full-length cell-free RNA sequencing, and here we show that nanopore sequencing can be leveraged to comprehensively characterize the vast, unannotated cell-free RNA transcriptome across healthy, precancerous, and cancerous disease states. By discovering and incorporating over 270,000 novel, unannotated cell-free RNAs into a custom transcriptome reference for quantification, feature selection, and machine learning, we demonstrate that cell-free RNAs enable highly accurate early detection of both precancerous high-grade dysplasia and esophageal adenocarcinoma, while providing systemic molecular insights into potential therapeutic targets and the progression from precancer to cancer.

## RESULTS

### LOCATE-seq RNA liquid biopsy platform for disease discovery

We developed a new RNA liquid biopsy platform technology called LOCATE-seq that harnesses <u>lo</u>ng-read sequencing of <u>c</u>ell-free RNA to <u>a</u>nnotate and quantify known and novel transcripts to <u>e</u>nable disease detection / monitoring, treatment selection, and drug discovery (Fig. 1a). We isolated cell-free RNA from the plasma of 16 healthy individuals, 12 patients diagnosed with precancerous Barrett’s esophagus with high-grade dysplasia, and 19 patients diagnosed with esophageal adenocarcinoma (n=13 stage I, n=5 stage II, n=1 stage III) (Table 1). We generated full-length cDNA libraries for long-read nanopore sequencing on the PromethION 48 sequencer. For novel transcript discovery^22^, we compiled all 47 nanopore sequencing libraries (40.7 million total reads), which revealed 276,346 previously unannotated transcripts that were not found in the GENCODE 39 annotation of the human transcriptome. Of these novel cfRNA transcripts, 270,679 occurred in intergenic regions.

**Fig. 1.**
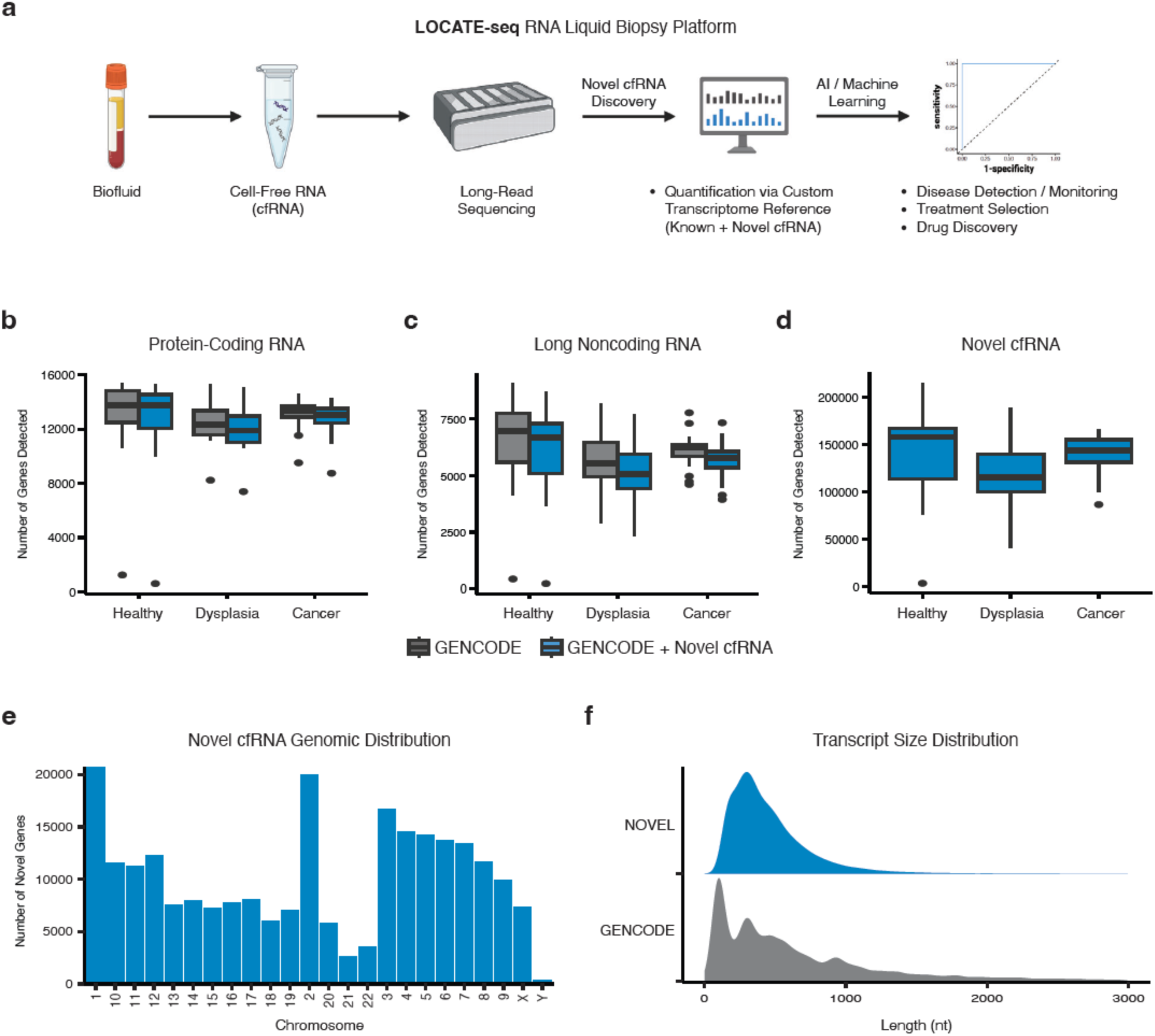
LOCATE-seq RNA liquid biopsy platform for disease discovery. a. LOCATE-seq schematic. b. Number of protein-coding RNAs detected across patient samples and transcriptome references. c. Number of long noncoding RNAs detected across patient samples and transcriptome references. d. Number of novel cfRNAs detected across patient samples. e. Genomic distribution of novel cfRNAs. f. Size distributions of novel and GENCODE cfRNAs.

**Table 1.** Patient plasma metadata.

| <u>Sample ID</u> | <u>Sex</u> | <u>Age</u> | <u>Histology</u> | <u>Cancer Stage</u> |
| --- | --- | --- | --- | --- |
| A1 | male | 76.2 | adenocarcinoma | I |
| A2 | male | 76.4 | adenocarcinoma | I |
| A3 | male | 68.7 | adenocarcinoma | I |
| A4 | male | 80.1 | adenocarcinoma | I |
| A5 | male | 61.8 | adenocarcinoma | II |
| A6 | male | 56.5 | adenocarcinoma | I |
| A7 | female | 82.1 | adenocarcinoma | I |
| A8 | female | 52.4 | adenocarcinoma | II |
| A9 | male | 55.5 | adenocarcinoma | I |
| A10 | female | 78.4 | adenocarcinoma | I |
| A11 | male | 59.7 | adenocarcinoma | II |
| A12 | female | 59.7 | adenocarcinoma | II |
| A13 | male | 51 | adenocarcinoma | I |
| A14 | female | 50.8 | adenocarcinoma | I |
| A15 | male | 55.5 | adenocarcinoma | I |
| A16 | female | 80.8 | adenocarcinoma | I |
| A17 | male | 75.3 | adenocarcinoma | II |
| A18 | male | 57.9 | adenocarcinoma | III |
| A19 | male | 83.2 | adenocarcinoma | I |
| B1 | male | 85.3 | high grade dysplasia |  |
| B2 | male | 61.4 | high grade dysplasia |  |
| B3 | male | 70.3 | high grade dysplasia |  |
| B4 | male | 64.5 | high grade dysplasia |  |
| B5 | male | 49 | high grade dysplasia |  |
| B6 | male | 68.4 | high grade dysplasia |  |
| B7 | male | 82.8 | high grade dysplasia |  |
| B8 | male | 67.9 | high grade dysplasia |  |
| B9 | male | 61.9 | high grade dysplasia |  |
| B10 | male | 88.4 | high grade dysplasia |  |
| B11 | male | 59.2 | high grade dysplasia |  |
| B12 | male | 42.6 | high grade dysplasia |  |
| C1 | male | 72 | control |  |
| C2 | male | 70 | control |  |
| C3 | male | 74 | control |  |
| C4 | male | 81 | control |  |
| C5 | male | 84 | control |  |
| C6 | female | 89 | control |  |
| C7 | male | 86 | control |  |
| C8 | male | 72 | control |  |
| C9 | male | 60 | control |  |
| C10 | male | 52 | control |  |
| C11 | female | 57 | control |  |
| C13 | female | 72 | control |  |
| C14 | female | 80 | control |  |
| C15 | male | 82 | control |  |
| C16 | male | 82 | control |  |

We next generated a custom transcriptome reference by combining our 276,346 novel cfRNA transcripts with all of the known transcripts in GENCODE 39, substantially expanding the known human cfRNA transcriptome. To more deeply profile our healthy and patient plasma samples, we then performed high-depth, short-read sequencing of our cfRNA libraries on the NovaSeq X sequencer (1.26 billion total reads) and mapped them to either the GENCODE transcriptome reference or our custom cfRNA transcriptome reference (GENCODE + Novel cfRNA) for quantification of both well-annotated GENCODE biotypes and the novel cfRNAs we discovered using long-read nanopore sequencing of full-length cfRNA. The median numbers of detected protein-coding genes were comparable whether we used the GENCODE transcriptome reference (healthy: 13,772, dysplasia: 12,342, cancer: 13,388) or our custom GENCODE + novel cfRNA transcriptome reference (healthy: 13,779, dysplasia: 11,892, cancer: 13,046) for cfRNA quantification (Fig. 1b). Median lncRNA genes detected were also comparable between GENCODE (healthy: 6,964, dysplasia: 5,540, cancer: 6,256) and GENCODE + novel cfRNA (healthy: 6,673, dysplasia: 5,074, cancer: 5,797) quantification (Fig. 1c). For novel cfRNAs, the number of detected genes was highest in healthy individuals (median: 157,821) when compared to dysplasia (median: 115,184) and cancer (median: 143,431) patients (Fig. 1d). Novel cfRNAs were distributed throughout the genome (Fig. 1e), with the majority of novel cfRNAs less than 1,000 nucleotides (nt) in length (Fig. 1f).

### Mitochondrial RNA enrichment in dysplasia and cancer

Quantification of cfRNA using the GENCODE transcriptome reference and subsequent differential expression (DE) analysis revealed significant enrichment of mitochondrial and protein-coding RNA in precancerous high-grade dysplasia (Fig. 2a), esophageal cancer (Fig. 2b), and both dysplasia and cancer cfRNA (Fig. 2c) when compared to healthy individuals. The mitochondrial cfRNA signal was the most significantly enriched molecular signature across all three DE analyses, indicating that mitochondrial cfRNAs that are strongly secreted at the precancerous state continue to persist in esophageal cancer patients. Protein-coding RNAs were the most commonly enriched across each of the DE analyses (Fig. 2d), while lncRNAs were the most commonly depleted across each of the comparisons (Fig. 2e). Significantly more mitochondrial genes were represented in the cfRNA of patients with precancerous dysplasia or cancer (Fig. 2f), suggesting that mitochondrial RNAs may serve as highly abundant biomarkers for both precancer and cancer early detection. Notably, a subset of dysplasia and cancer patients exhibited substantially higher levels of mitochondrial RNA in their cfRNA transcriptomes, while other patients showed more moderate levels of mitochondrial RNA enrichment (Fig. 2g).

**Fig. 2.**
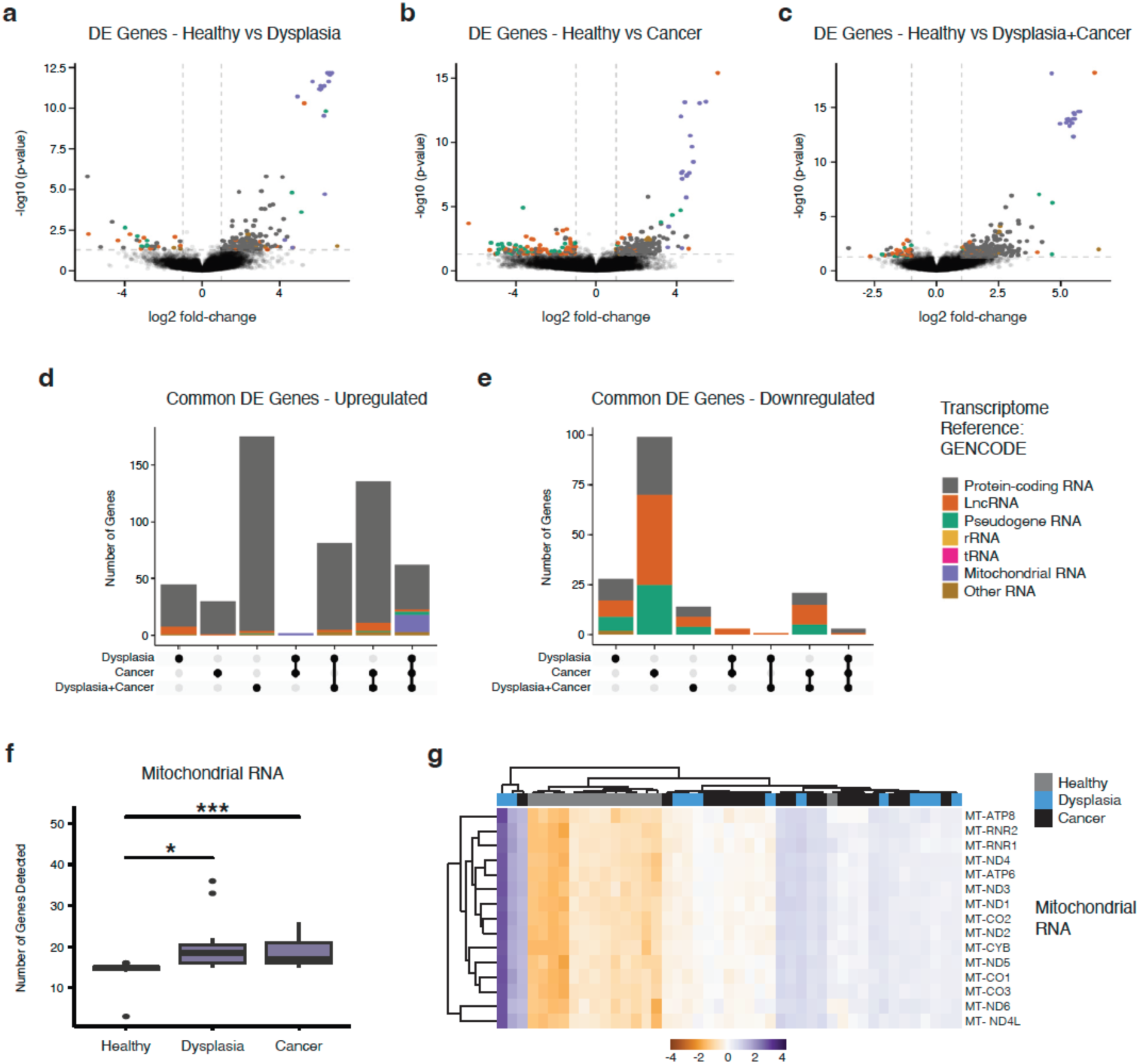
Mitochondrial RNA enrichment in dysplasia and cancer. a. Differential expression (DE) analysis volcano plot of healthy vs. dysplasia cfRNA. b. DE analysis volcano plot of healthy vs. cancer cfRNA. c. DE analysis volcano plot of healthy vs dysplasia and cancer cfRNA. d. UpSet plot of common upregulated DE genes. e. UpSet plot of common downregulated DE genes. f. Box plot of number of detected mitochondrial genes. g. Hierarchical clustering of mitochondrial RNA expression.

### Differential abundance of novel cfRNA in dysplasia and cancer

We next used our custom transcriptome reference containing all of our novel cfRNA transcripts (GENCODE + novel cfRNA) for quantification and DE analysis in precancerous high-grade dysplasia when compared to healthy (Fig. 3a), esophageal cancer when compared to healthy (Fig. 3b), and when comparing healthy to both dysplasia and cancer cfRNA (Fig. 3c). Protein-coding RNAs were again the most commonly enriched in disease across each of the comparisons (Fig. 3d), while novel cfRNAs were significantly depleted in dysplasia or cancer cfRNA (Fig. 3e). We also found a cluster of 20 novel cfRNAs that exhibited higher overall expression across both dysplasia and cancer patients (Fig. 3f).

**Fig. 3.**
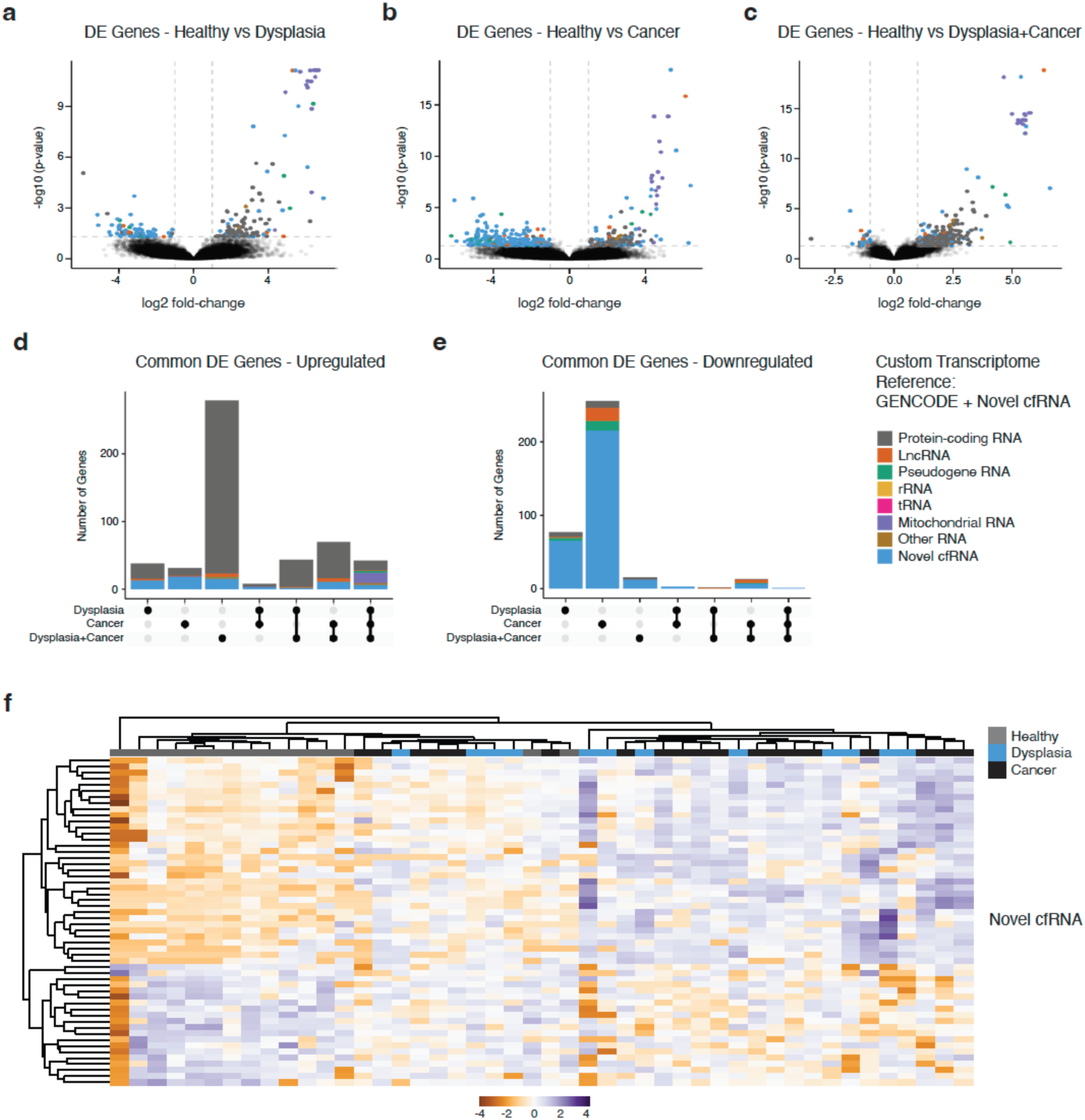
Differential abundance of novel cfRNA in dysplasia and cancer. a. Differential expression (DE) analysis volcano plot of healthy vs. dysplasia cfRNA. b. DE analysis volcano plot of healthy vs. cancer cfRNA. c. DE analysis volcano plot of healthy vs dysplasia and cancer cfRNA. d. UpSet plot of common upregulated DE genes. e. UpSet plot of common downregulated DE genes. f. Hierarchical clustering of novel cfRNA expression.

### Novel cfRNA features enable accurate disease classification

When we examined novel cfRNAs that were unique to each individual, we observed a significant increase in unique novel cfRNAs in esophageal cancer patients (Fig. 4a), with a few patients expressing over 1,000 unique novel cfRNAs (Fig. 4b). Using a feature set that contained both well-annotated GENCODE transcripts and our novel cfRNA transcripts, we trained machine learning models (logistic regression) with L1 regularization using the significantly DE cfRNAs as input features (p-adjusted < 0.05). Our models segregated both high-grade dysplasia (Fig. 4c) and esophageal cancer (Fig. 4d) from healthy samples with perfect sensitivity (100%) and specificity (100%) using a limited number of features, highlighting the utility of our approach for both precancer and cancer early detection using noninvasive RNA liquid biopsy technology.

**Fig. 4.**
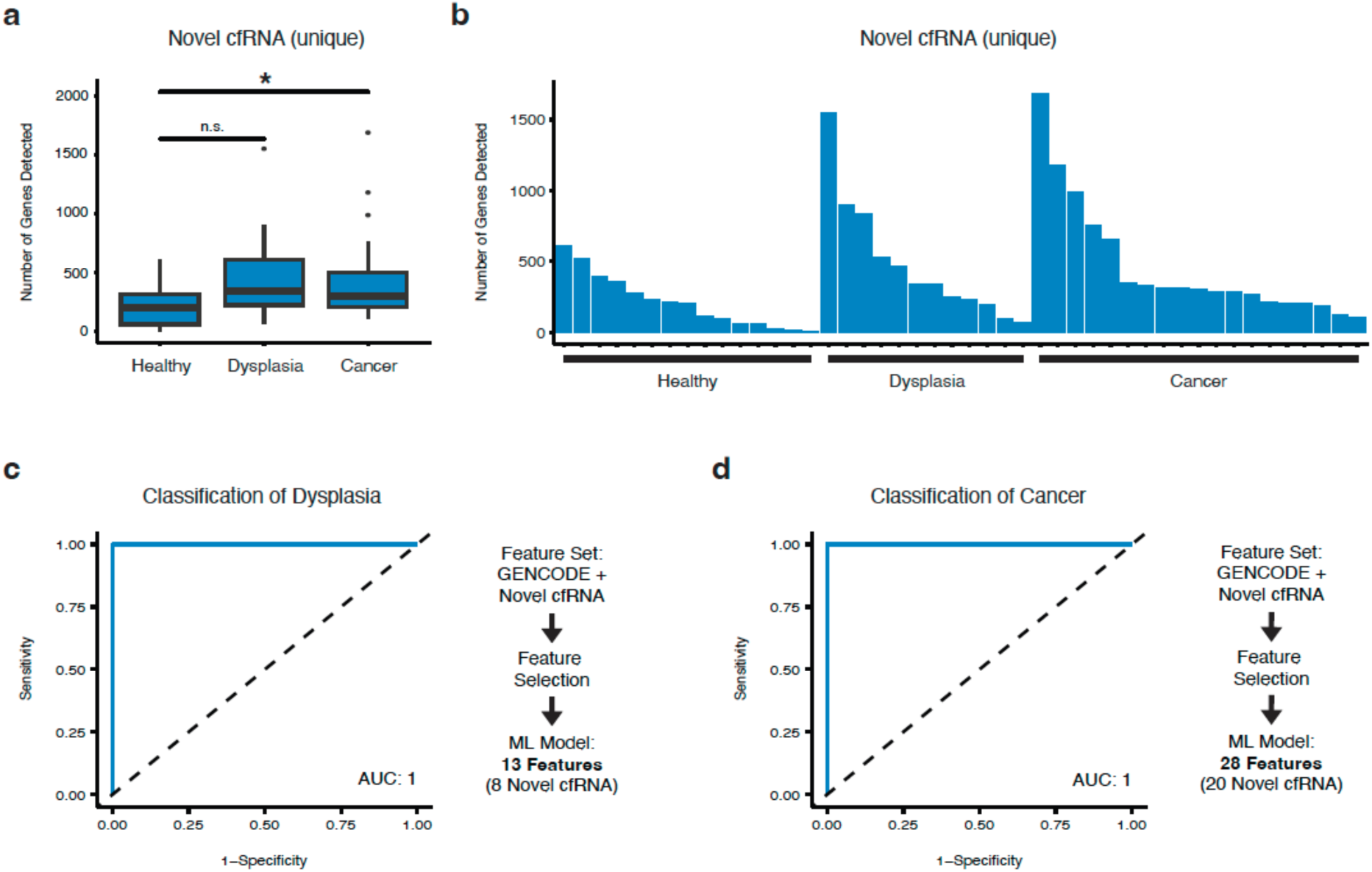
Novel cfRNA features enable accurate disease classification. a. Box plot of number of detected unique novel cfRNAs. b. Bar plot of number of detected unique novel cfRNAs in each healthy individual or patient. c. Receiver operating characteristic curve for logistic regression model trained using known and novel cfRNA features for classification of high-grade dysplasia. d. Receiver operating characteristic curve for logistic regression model trained using known and novel cfRNA features for classification of early-stage esophageal cancer.

### Dysregulated pathways as potential therapeutic targets in dysplasia and cancer

We next characterized common gene sets and pathways that were significantly dysregulated in both high-grade dysplasia and early-stage esophageal cancer patients to identify potential therapeutic targets. We performed gene seat enrichment analysis (GSEA) and found that “MYC targets” and the mitochondrial process of “oxidative phosphorylation” were the top 2 enriched gene sets across both dysplasia and early-stage cancer (Fig. 5a). “PI3K AKT MTOR signaling” and “MTORC1 signaling” were also among the top 10 enriched gene sets in both dysplasia and cancer (Fig. 5a). The plasma cfRNA levels of biomarkers such as human epidermal growth factor-2 (HER2) remained relatively constant between healthy individuals and patients with dysplasia or cancer (Fig. 5b), but the cfRNA levels of immunotherapy biomarkers such as the immune checkpoint genes CTLA-4 (Fig. 5c) and LAG-3 (Fig. 5d) increased in the plasma of Barrett’s esophagus patients with high-grade dysplasia. Plasma cfRNA levels for immune checkpoint genes PD-L1 (Fig. 5e) and TIGIT (Fig. 5f) and the p53 regulator MDM2 (Fig. 5g) were relatively consistent across all of the sample cohorts, although a few cancer patients exhibited much higher levels of these biomarkers.

**Fig. 5.**
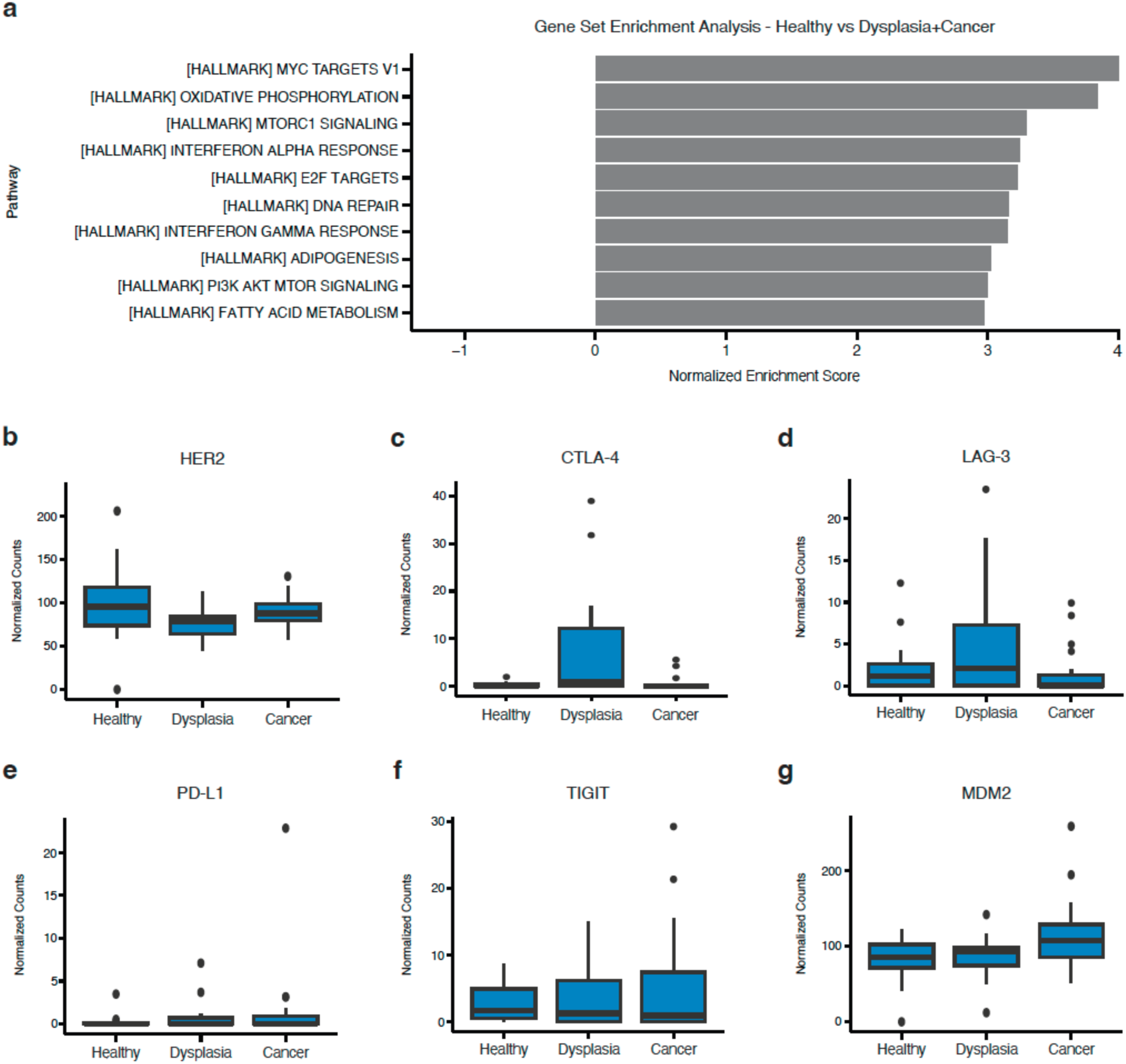
Dysregulated pathways as potential therapeutic targets in dysplasia and cancer. a. Gene set enrichment analysis for pathways enriched in dysplasia patient plasma (top 10 most significantly enriched pathways). b-g. Box plots of normalized counts for HER2, CTLA-4, LAG-3, PD-L1, TIGIT, and MDM2 cfRNA.

## DISCUSSION

Here we show that the comprehensive profiling of cfRNA using full-length nanopore sequencing enables the discovery of 276,346 unannotated, novel cfRNA transcripts, many of which are differentially expressed in disease and can be leveraged as biomarkers for precancer and cancer early detection with high sensitivity and specificity using machine learning. Our findings reveal that our understanding of the true complexity of cfRNA transcriptomes in the context of health and disease is still nascent, and by using single-molecule, long-read nanopore sequencing of full-length cfRNA, we reveal the dynamic diversity of cfRNA transcriptomes in healthy individuals and patients with precancerous high-grade dysplasia or esophageal adenocarcinoma. Our LOCATE-seq RNA liquid biopsy platform technology allows us to profile and harness this rich diversity of cfRNA for novel biomarker discovery and highly accurate disease classification.

Our study also reveals that mitochondrial RNA are highly enriched in the cfRNA in both precancerous dysplasia and early-stage cancer patient plasma, highlighting their utility as biomarkers of disease and underscoring their role in providing potential insights into metabolic changes and disease biology. A recent study using the PC3 prostate cancer cell line also showed that mitochondrial RNA are enriched in large oncosomes, suggesting that mitochondrial RNA may serve as circulating biomarkers for various types of cancers^23^. Increased levels of mitochondria-derived cfRNA and the enrichment of gene sets such as oxidative phosphorylation in dysplasia and cancer also suggest potential therapeutic strategies for disease treatment using drugs that target this metabolic pathway. Our findings also uncover additional pathways for potential therapeutic intervention, including enrichment of the PI3K AKT MTOR signaling pathway in both dysplasia and cancer, and increased levels of immune checkpoint pathway genes such as CTLA-4 and LAG-3 in precancerous dysplasia, suggesting the potential utility of immunotherapy for esophageal cancer prevention.

Taken together, our study provides proof-of-concept that long-read nanopore sequencing of cfRNA reveals a rich diversity of previously unknown cfRNA transcripts that can serve as new biomarkers and novel features for precancer and cancer early detection using machine learning / artificial intelligence. Further validation using plasma samples from independent patient cohorts will be important for validating and testing the performance of our models. Our study also demonstrates the advantages of comprehensively profiling the cfRNA transcriptome for drug target discovery and for informing potential therapeutic strategies for cancer treatment and prevention.

## METHODS

### Cell-free RNA isolation

Blood plasma from patients were collected from Cambridge University Hospital. Healthy donors were identified as those without disease by BioIVT sample metadata. Plasma samples were initially run through a 0.8µm filter to remove cellular debris. Cell-free RNA was isolated from the filtered plasma using the Norgen Biotek Plasma/Serum Circulating and Exosomal RNA Purification Kit (slurry format). RNA was eluted into 50 µl and stored at –80C prior to cDNA synthesis.

### cDNA synthesis

10 ng of input RNA was used for the Takara SMART-Seq HT plus kit. cDNA was prepared as specified by the manufacturer’s protocol and amplified for 24 cycles. Quantification of the resulting cDNA was done using both a Nanodrop spectrophotometer, as well as a Qubit Fluorometer. The size distribution of the cDNA was verified using an Agilent Bioanalyzer prior to sequencing library preparation.

### Long-read sequencing

Barcoded nanopore sequencing libraries were created using a custom low-input protocol for the Oxford Nanopore Technologies SQK-NBD-114.24 and LSK-114 kits. Briefly, cDNA was end-repaired and A-tailed as specified, but incubated for 30 minutes, followed by a 30-minute deactivation. Barcodes from the SQK-NBD-114.24 kit were incubated at 20 C for 4.5 hours. Ligation was terminated by the addition of 2ul of EDTA to each reaction. Barcoded samples were pooled and subject to a 1.8x Ampure Bead cleanup, and then used as input for the LSK-114 protocol. Here, we modified all bead cleanups to use a 1.8x ratio and performed all bead washes at 37 C. Adapter ligation was performed for 20 minutes to promote ligation yield. Short fragment buffer was used to capture the full gamut of cDNA lengths. The final library was eluted into 25ul of elution buffer and loaded onto an R10.4 Promethion flow cell. POD5s were basecalled and demultiplexed using Dorado version 0.5.3 using the SUPv4.3.0 basecalling model. Reads that failed demultiplexing were re-demultiplexed using a custom python script that exact matches barcodes and later combined with their respective FASTQs.

### Short-read sequencing

Illumina libraries were prepared using full-length cDNA generated from the isolated plasma RNA of healthy, Barrett’s esophagus, and esophageal cancer samples using the Takara SMART-Seq HT plus kit and sequenced on an Illumina NovaSeq (2×150 bp reads).

### Novel transcript discovery

Isoquant version 3.3.1 with the Human reference genome build HG38 and the GENCODE version 39 comprehensive transcript annotation set was used to identify novel transcripts against all FASTQs in one batch with the following parameters:

-d nanopore, –stranded none, --report_novel_unspliced true, –matching_strategy loose, --gene_quantification all, --transcript_quantification all.

Novel transcripts were merged with the GENCODE annotations. Assembly quality was assessed using SQANTI version 3 with default parameters.

### RNA-seq quality and quantification

RNA-seq short reads (FASTQ) were trimmed using FastP (v0.23.4), assessed for quality using FastQC (v0.12.1), and quantified using Salmon (v1.10.1). Quantification was run with the following parameters:

--gcBias, --seqBias, --validateMappings, --recoverOrphans, --rangeFactorizationBins 4 To reduce sequences biases, enable selective alignments, rescue reads with an unmapped pair, and improve quantification accuracy. The salmon mappings were performed against a concatenation of:

1. the HG38 human genome reference annotation (v39) from the GENCODE consortium, and the RepeatMasker track from the University of California Santa Cruz genome browser
2. The above, and an annotation of novel genes identified via IsoQuant (3.3.1)

This assembled a transcriptome annotated with the corresponding ENSEMBL transcript IDs and novel gene IDs from IsoQuant. Transcript counts were aggregated to the gene level for GENCODE genes.

### Short-read analysis

Short-read transcripts were imported into R (v4.3.3) using tximport (v1.22.0). Biotypes were assigned to genes using their respective GENCODE biotypes. IsoQuant annotated novel genes were assigned the ‘Novel’ biotype. Count normalization and differential expression analyses were performed using DESeq2 (v1.42.0). Significant differential expression was limited to genes with a Benjamini-Hochberg adjusted p-value < 0.05, and an absolute value log2 fold-change > 1.

### Modeling

Logistic regression models were train in R using the glmnet package (v4.1), with the identified differentially expressed genes as input features. Models were trained with respect to the sample age and sex (~condition+age+sex). One sample was missing the age information in the metadata, and the average age of samples with the same histology and sex was used. Optimal lambda values for the logistic regression model were identified using 5-fold cross validation, and the value 1 standard error away from the minimum cross-validated value of lambda was selected for final model training. The final model was trained utilizing the entire dataset, and further feature reduction was performed using LASSO regression.

## Data availability

Nanopore long read sequencing data have been deposited at the NCBI Gene Expression Omnibus repository under accession number GSE271389 and Illumina short read sequencing data under accession number GSE271390.

## ACKNOWLEDGEMENTS

We thank Hannah Coles, Adam Freeman, and the OCCAMS consortium for providing plasma samples and anonymized data, and infrastructure was supported by Cancer Research UK (CRUK) on behalf of the OCCAMS consortium. This work was supported by Cancer Research UK [A15874, A22720, A22131] and the Medical Research Council [MR/W014122/1] to R.C.F. with additional support by the UK National Institute for Health Research (NIHR) Cambridge Biomedical Research Centre (NIHR203312), and the American Cancer Society (https://doi.org/10.53354/pc.gr.158353) and philanthropic support to D.H.K. V.P. was supported by the Tobacco-Related Disease Research Program (T32DT4904), A.H. was supported by the UCSC Committee of Research Large Grant Program (09507-444096), S.V.M. was supported by the California Institute for Regenerative Medicine (EDUC4-12759), and D.H.K. was supported by a Research Scholar Grant (RSG-22-099-01-CDP) from the American Cancer Society.

## AUTHOR CONTRIBUTIONS

D.H.K. conceived and designed the study, V.P. and A.H. performed computational analyses and generated figures, S.V.M. and C.M. performed experiments, J.G., K.M., and R.F. provided materials, D.H.K. supervised research, and D.H.K., V.P., and A.H. wrote the paper with input from the authors.

